# Contextual control of conditioned pain tolerance and endogenous analgesic systems: Evidence for sex-based differences in endogenous opioid engagement

**DOI:** 10.1101/2021.12.02.470965

**Authors:** Sydney Trask, Jeffrey S. Mogil, Fred J. Helmstetter, Cheryl L. Stucky, Katelyn E. Sadler

**Affiliations:** Department of Psychological Sciences, Purdue University; Departments of Psychology and Anesthesia, Alan Edwards Centre for Research on Pain, McGill University; Department of Psychology, University of Wisconsin-Milwaukee; Department of Cell Biology, Neurobiology and Anatomy, Medical College of Wisconsin

**Author notes:** Contact: Katelyn E. Sadler, Address: 8701 Watertown Plank Road, Milwaukee WI, 53226.

## Abstract

The mechanisms underlying the transition from acute to chronic pain are unclear but may involve the persistence or strengthening of pain memories acquired in part through associative learning. Contextual cues, which comprise the surrounding environment where events occur, were recently described as a critical regulator of pain memory; both rodents and humans exhibit increased pain sensitivity in environments recently associated with a single painful experience. It is unknown, however, how repeated exposure to an acute painful unconditioned stimulus in a distinct context modifies pain sensitivity or the expectation of pain in that environment. To answer this question, we conditioned mice to associate distinct contexts with either repeated administration of a mild visceral pain stimulus (intraperitoneal injection of acetic acid) or vehicle injection over the course of three days. On the final day of experiments animals received either an acid injection or vehicle injection prior to being placed into both contexts. In this way, contextual control of pain sensitivity and pain expectation could be tested respectively. Both male and female mice developed context-dependent conditional pain tolerance, a phenomenon mediated by endogenous opioid signaling. However, when expecting the presentation of a painful stimulus in a given context, males exhibited conditional hypersensitivity whereas females exhibited endogenous opioid-mediated conditional analgesia. Successful determination of the brain circuits involved in this sexually dimorphic anticipatory response may allow for the manipulation of pain memories, which may contribute to the development of chronic pain states.

## Introduction

Chronic pain development may involve the generalization and strengthening of acute pain memories (Moseley & Vlaeyen, 2015). Neuronal encoding of painful stimuli, known as nociception, is an evolutionarily conserved, unconditional response (UCR) generated upon exposure to a tissue-damaging unconditional stimulus (UCS). When pain persists, subjects are either repeatedly presented with or chronically experiencing this UCS, providing opportunity for various external cues or conditional stimuli (CS) to become associated with the UCS. Pairings of the painful UCS with external cues can eventually lead to the development of a conditional response (CR) elicited by the conditional stimulus alone. In this way, nociceptive signaling and resulting pain behaviors may be modulated by previously biologically irrelevant stimuli.

Pavlov (1927) was the first to describe the process through which a biologically irrelevant stimulus could come to exert control over physiological processes in a way that allows the organism to prepare for the upcoming disruption of homeostasis produced by the unconditional stimulus. In addition to auditory stimuli like those used in some of Pavlov’s experiments, the immediate environment or “context” may control these preparatory responses as was demonstrated in a series of experiments performed by Siegel and colleagues (1982). Over the course of 15 days, rats were treated with escalating doses of the mu-opioid receptor agonist, heroin, in one distinct physical environment. After training, animals received a lethal dose of heroin (96.4% mortality rate in heroin-naïve rats) in the same environment where training occurred, or in a separate, novel environment. When administered in a shifted context, this lethal dose of heroin killed 64.3% of animals; heroin administration in the familiar context decreased the death rate by ∼50% such that only 32.4% of animals died in response to administration of the lethal dose. Similar to the conditional response first reported by Pavlov (1927), this work suggested that the familiar environment served as a “conditional stimulus” (CS) that predicted upcoming heroin administration, thereby initiating internal processes that mimic those associated with drug tolerance. This phenomenon has since been termed a “conditioned compensatory response” as exposure to the CS begins the process of compensating for the upcoming drug effect (see also Siegel, 1977). Here we used a similar experimental paradigm to determine if repeated exposure to an acute painful stimulus in a given context affects sensitivity to that stimulus. In other words, do animals exhibit conditioned compensatory responses to painful stimuli? And if they do, what is the physiological basis of that response?

Context has also been well-documented to exert control over learned behaviors related to fear and anxiety (Bouton & Bolles, 1979) as well as instrumental and choice behaviors (Trask et al., 2014). Recently, context was found to influence pain behaviors in male mice and humans after a single conditioning trial (Martin et al., 2019). Both male mice and men exhibited stress-induced hyperalgesia if tested in an environment previously associated with an unconditional painful stimulus (Martin et al., 2019). Neither female mice nor women exhibited conditional hypersensitivity or context-dependent increases in circulating stress hormones in these experiments, suggesting that context did not impact nociceptive signaling in females as it did in males. However, an alternative interpretation of these data may be that contextual cues engaged compensatory endogenous analgesic systems in females, even in the absence of ongoing pain, such that their pain behaviors pre- and post-conditioning were indistinguishable. This hypothesis is appealing given the potential therapeutic implications that could follow if environment is capable of modulating endogenous analgesic mechanisms. To date, the few studies that have examined environmental influence on conditional analgesia have primarily done so in fear conditioning paradigms. In general, those studies found that following fear conditioning in which a conditional stimulus predicts an aversive shock outcome, animals exhibited increased pain tolerance (*i*.*e*., conditional hypoalgesia) in the environment, or context, where fear conditioning occurred (Fanselow & Helmstetter, 1988; Helmstetter & Fanselow, 1987; Watkins et al., 1993). Although the conditional fear responses exhibited following this associative learning have been extensively studied, the potential compensatory analgesic responses produced by context exposure have received little to no attention outside of these paradigms. We therefore designed a novel set of experiments to determine if context could be used as a conditional stimulus to engage endogenous analgesic mechanisms and thus increase pain tolerance, in the absence of overt fear conditioning or ongoing, continuous pain.

## Methods

### Animals

Equally sized cohorts of male and female C57BL/6 mice aged 8-12 weeks were used in all experiments. Mice were either bred in house or purchased from The Jackson Laboratory (Bar Harbor, ME). Purchased animals acclimated to the housing facility for >7 days before experimental use. Purchased animals and animals bred in-house were not intermixed, but rather used in independent experiments. Animals were randomized to treatment group. All protocols were in accordance with National Institute of Health guidelines and were approved by the Institutional Animal Care and Use Committee at the Medical College of Wisconsin (Milwaukee, WI; protocol #0383). We are not the first to use repeated acetic acid injections as painful stimuli. This procedure was previously used in Stevenson et al. (2006); approximately 4 hr after acetic acid injection, animals no longer exhibited acetic acid-induced feeding suppression in those studies. Similarly, no animal in our study lost more than 15% of their body weight over the course of the experiments. Therefore no animals were excluded from these studies as dictated by approved IACUC endpoints.

### Conditioning paradigms

All experiments used two distinct contexts, counterbalanced as Context A and B for all animals. The first context was a 10 × 10 × 15-cm^3^ Plexiglas chamber. Two walls of the chamber were solid black Plexiglas, and a horizontal white and black striped pattern covered the remaining two walls. These chambers were cleaned with C-DOX, a chlorine-based disinfectant. The second chamber was a similarly sized Plexiglas chamber, consisting of four black walls, and cleaned with 70% ethanol. Both contexts were placed onto a raised 0.7-cm^2^ wire platform to allow for video recordings and mechanical sensitivity testing. Dilute acetic acid (0.1, 0.3, 0.5, or 0.9%) was used as the unconditioned stimulus in all experiments. Animals received an intraperitoneal (IP) injection (10 ml/kg body weight) of acetic acid (purchased from Sigma Aldrich; diluted in phosphate buffered saline, PBS, pH 7.4) or PBS vehicle according to the experimental details described below. The 0.9% dose of acetic acid was used as the final unconditional stimulus since it has previously been shown to elicit conditioned pain hypersensitivity in male mice (Martin 2019).

### Within-subject assessment of conditioned pain tolerance

On days 1, 2, and 3, animals received an IP injection of acetic acid immediately before being placed into one of the two contexts. In the escalating dose experiment (Figure 2), the concentrations were 0.1%, 0.3% and 0.5% acetic acid on days 1, 2, and 3 respectively. For the consistent dose experiments (Figure 3), a 0.9% acetic acid injection was given on each of days 1, 2, and 3. Animals received an equivalent volume injection of PBS immediately before being placed into the second of the two contexts. Contexts and time of injection (i.e., morning or afternoon) were counterbalanced across all animals and context pairings were separated by 3 h for each animal. On day 4, animals received an IP injection of 0.9% acetic acid immediately before being placed into Context A. A second IP injection of 0.9% acetic acid was administered 3 h later immediately before placing the animal into Context B. Pain behavior testing was completed after being placed into either context on days 1 and 4, as described below.

### Between-subject assessment of conditioned pain tolerance

Between-subjects experiments were similar to the within-subject experiments, with the exception that each animal was only exposed to one context. On days 1, 2, and 3, half of all animals received an IP injection of 0.9% acetic acid immediately before being placed into one of the two conditioning chambers described above; chambers were counterbalanced between groups and animals were only exposed to one of the contexts. The remaining half of the animals received an IP injection of PBS before being placed into the chamber. On day 4, all animals received an IP injection of 0.9% acetic acid immediately before being placed into their training context. Pain behavior testing was completed after being placed into the context on days 1 and 4 as described below.

### Pain behavioral measures

Animals were moved to the behavior testing room at approximately 07:00 h each morning. Overhead lights were on throughout the entirety of the habituation and testing period. Animals remained in their home cages in the behavior room for no less than 1 h prior to the first injection each day. Immediately following acetic acid or vehicle injection, animals were placed into Context A or Context B. Mechanical sensitivity was assessed in each animal 45-60 min following injection using von Frey filaments. Calibrated monofilaments were delivered through the wire testing platform and applied to the plantar surface of each hindpaw following the up-down method (Dixon 1965); the 50% withdrawal threshold of each paw was calculated then averaged between paws (Dixon 1980, Chaplan et al., 1994). Toe flaring was not considered a withdrawal.

### Naloxone treatment

Naloxone hydrochloride was purchased from Sigma Aldrich and dissolved in PBS, pH 7.4. Naloxone or an equivalent volume of PBS vehicle was administered via IP injection to result in a final dose of 10 mg/kg. This dose has previously been shown to block the contribution of endogenous opioid signaling in the tail-withdrawal test (Rosen et al., 2019) and reduce spontaneous pain behaviors initiated by complete Freund’s adjuvant injection (Lee et al., 2021).

### Corticosterone ELISA

Immediately following von Frey mechanical sensitivity testing (*i*.*e*., 60 min following acetic acid or vehicle injection on day 4; Figure 3), animals were removed from the testing apparatus, placed into a clean shoebox cage, and transferred (<1 min) to a neighboring necropsy room. Trunk blood was obtained from isoflurane anesthetized mice via cardiac puncture, transferred to a chilled tube containing 3.5% sodium citrate, then centrifuged at 1500 rpm, 4 °C for 15 min. Plasma was collected and stored at -80 °C until analysis. Plasma concentrations of corticosterone were measured using the Corticosterone Enzyme Immunoassay kit (Arbor Assay’s DetectX) as previously described (Long et al., 2016).

### Data reporting and analysis

Data presented in this manuscript are those collected during the first performance of each experiment. Results for all animals enrolled in each experiment are reported; no outliers were encountered. All data were analyzed using repeated measures ANOVAs with α=0.05 in SPSS v.28. Planned comparisons were used following ANOVAs to determine between- and within-group differences following significant interactions or main effects. Estimates of effect size were calculated using a partial eta-squared. Power analyses were used to determine appropriate sample sizes. Means (μ), standard deviations (σ), and expected effect sizes were obtained from preliminary studies and data of a similar nature previously published by the Stucky Lab. These values were used to calculate the sample sizes needed to achieve a significance level (α) of 0.05 and statistical power (1-β) of >0.8.

## Results

In chronic pain conditions and models, subjects are repeatedly or continuously presented with a noxious UCS. In these studies, we used associative learning paradigms that repeatedly coupled a painful UCS (*i*.*e*., acetic acid) with a unique environment to determine if pain memories affect (1) pain sensitivity (UCR), and (2) pain expectation (CR) **(Fig. 1)**.

**Figure 1:**
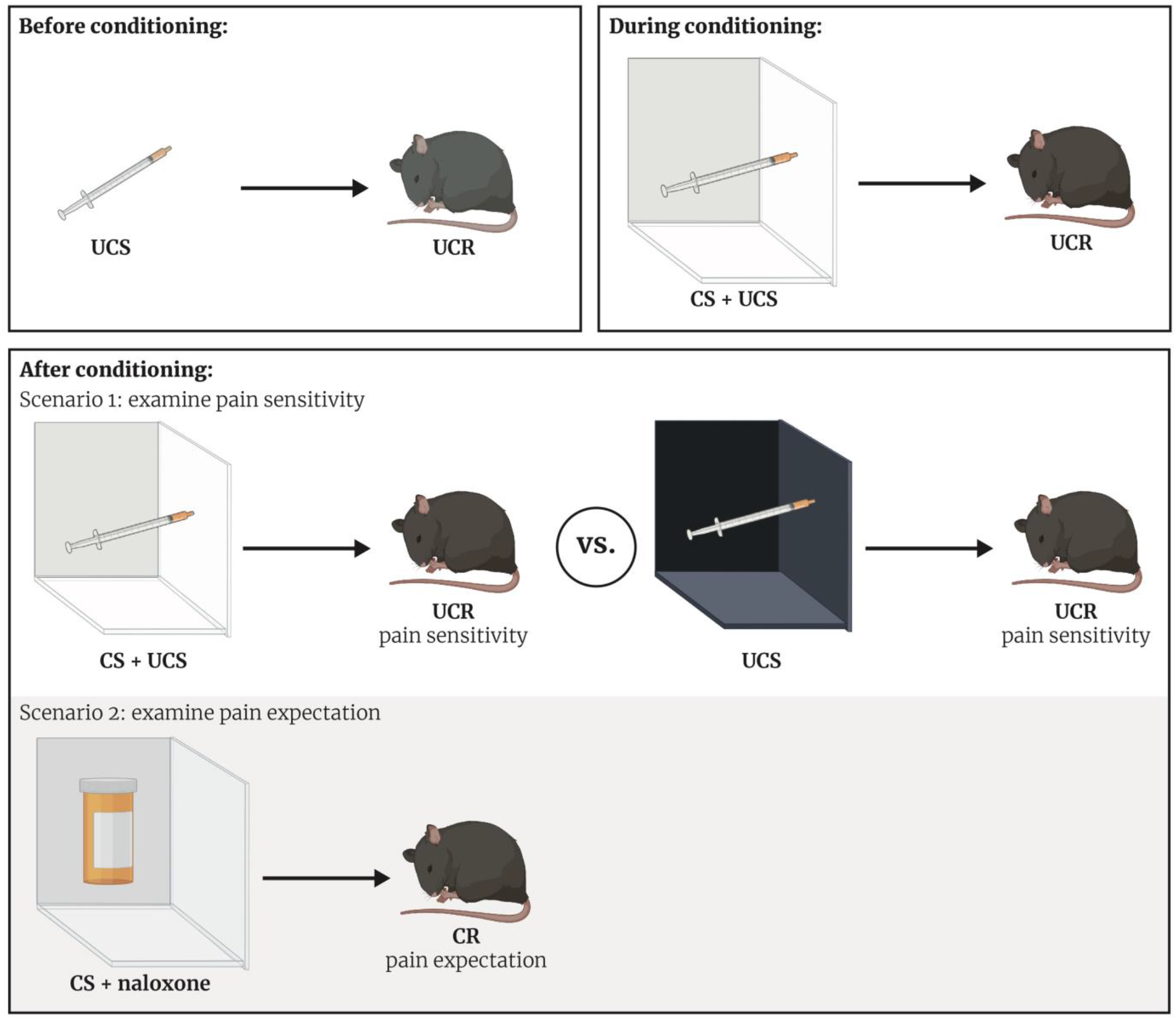
Using associative learning to investigate contextual control over pain sensitivity and pain expectation. Pavlovian conditioning was used to determine if context could come to exert control over different aspects of pain behaviors. Prior to conditioning, the unconditional stimulus (UCS; 0.9% IP acetic acid injection) induced pain behaviors (unconditional response, UCR; hindpaw mechanical hypersensitivity) in mice. During conditioning, the UCS was presented immediately before animals were placed into a unique training environment (conditional stimulus, CS; behavior chambers with unique wall patterns and odors), such that the UCR occurred in the CS. After conditioning, two different experiments were performed to assess how environment influences both pain sensitivity (UCR) and pain expectation (conditional response, CR). In the first experiment, the UCS was presented in both the CS and a novel environment that had different visual, and olfactory cues. The UCR was measured in each context and resulting differences determined if environment affected pain sensitivity. In the second experiment, animals were treated with naloxone and placed into the CS after multiple presentations of the UCS + CS to determine if endogenous opioid signaling influences conditional behavioral responses.

### Female mice develop context-dependent pain tolerance after training with ascending doses of acetic acid

To first determine if context can exert conditional control over pain sensitivity, we designed a within-subject, 3-day conditioning paradigm in which animals were trained to associate one unique context with escalating doses of IP acetic acid and a second context with IP vehicle administration (Fig. 2A). Prior to being placed into either chamber on the fourth day of the paradigm, animals were injected with a dose of acetic acid that was higher than the doses previously administered in the acid-trained context. The primary pain behavior measurement in these experiments was mechanical sensitivity of the hindpaw. Although the hindpaw tissue is not directly injured by the UCS, the sensitivity of the hindpaw may be altered by ongoing visceral pain through classical referred pain mechanisms (*i*.*e*., sensitization of spinal cord projection neurons receiving information from both the foot and the viscera), or through the recently described ‘anti-diffuse noxious inhibitory control (anti-DNIC)’ phenomenon in which pain in one region of the body increases pain in another anatomically distinct region (Tansley et al., 2019). Hindpaw sensitivity was measured in both contexts on day 1 and day 4 of the experiment, 45 min following acetic acid injection. A 2 (Sex: Male, Female) x 2 (Context: Vehicle-Paired, Acetic Acid-Paired) x 2 (Day: 1, 4) ANOVA revealed a main effect of day, *F*_(1, 14)_ = 19.08, MSE = 1.71, *p* = 0.001, ηp^2^ = 0.58, a day by sex interaction, *F*_(1, 14)_ = 15.51, MSE = 1.71, *p* = 0.009, ηp2 = .39, a context by day interaction, *F*_(1, 14)_ = 13.75, MSE = 1.81, *p* = 0.002, ηp^2^ = 0.50, and a three-way interaction, *F*_(1, 14)_ = 23.84, MSE = 1.81, *p* < 0.001, ηp^2^ = 0.63. There was no main effect of context, sex, nor an interaction between the two, *F*s < 1. Planned comparisons were further conducted to assess differences both between and within groups. Male mice **(Fig. 2B)** exhibited similar levels of hindpaw mechanical sensitivity in both the acid- and vehicle-paired contexts on day 1 (*p* = 0.91) and day 4 (*p* = 0.29), suggesting that associative learning did not enable environmental control of pain sensitivity in males. Alternatively, hindpaw sensitivity of female mice varied greatly between chambers on day 1 and day 4 of the paradigm **(Fig. 2C)**. On day 1, female mice exhibited mechanical hypersensitivity (*i*.*e*., ‘anti-DNIC’) in the acid-paired context relative to the vehicle chamber (*p* < 0.001). However, on day 4, female hindpaw sensitivity was greater in the vehicle-paired context than the acid-paired context (*p* = 0.001), despite the fact that animals received IP injection of 0.9% acetic acid prior to being placed into either chamber.

**Figure 2:**
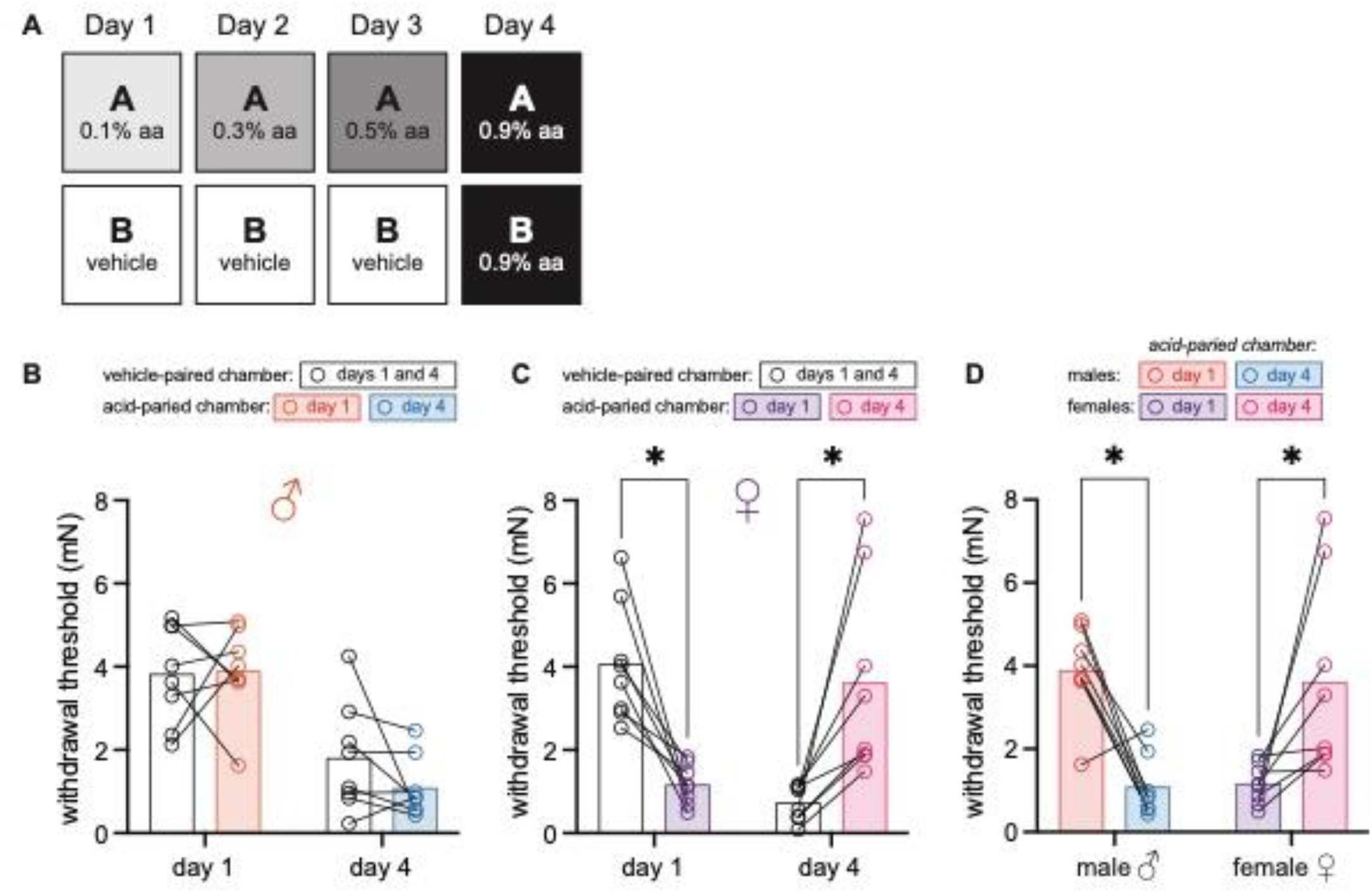
Female mice develop context-dependent pain tolerance after training with ascending doses of acetic acid. **A)** Experimental design depicting the within-subject procedure. In daily sessions, mice were given ascending doses of acetic acid in one physical chamber, Context A, and vehicle injections in a separate physical chamber, Context B. On the final day, mice received an injection of 0.9% acetic acid solution in each context. Hindpaw sensitivity was measured 30 min following injection on day 1 and day 4. **B)** von Frey withdrawal thresholds of male mice (n=8) on day 1 and day 4 of the paradigm. Acetic acid injection on day 1 (0.1%) had no effect on hindpaw mechanical sensitivity. Hindpaw mechanical sensitivity was similar in both contexts following 0.9% acetic acid injection on day 4. **C)** von Frey withdrawal thresholds of female mice (n=8) on day 1 and day 4 of the paradigm. Acetic acid injection on day 1 (0.1%) induced hindpaw mechanical hypersensitivity (‘anti-DNIC’). Hindpaw mechanical sensitivity differed between the contexts following 0.9% acetic acid injection on day 4, however; females exhibited contextually mediated pain tolerance (i.e., less pain sensitivity) in the acid-paired chamber relative to the vehicle-paired chamber on day 4. **D)** von Frey withdrawal thresholds of male and female mice replotted to highlight results in the acetic-acid paired chamber. Male mice exhibit anti-DNIC on day 4 relative to day 1. Female mice exhibit less hindpaw sensitivity following 0.9% acetic acid injection on day 4 than they did following 0.1% acetic acid injection on day 1 suggesting the development of conditional pain tolerance.

The individual sexes exhibited similar levels of hindpaw sensitivity in the vehicle-paired chamber on day 1 (*p* = 0.72), but displayed both quantitative and qualitative differences in hindpaw sensitivity following the various acetic acid injections. Following acid injection on day 1, female hindpaw mechanical sensitivity was higher than that observed in males (*p* < 0.001; **Fig, 2D)**. Similarly, female hindpaw sensitivity was greater than male hindpaw sensitivity following an IP injection of 0.9% acetic acid and placement into the vehicle-paired chamber on day 4 (*p* = 0.045), despite the fact that both male and female mice exhibited higher mechanical sensitivity in this chamber on day 4 compared to day 1 (males: *p* = 0.003; females: *p* < 0.001). However, when injected with acid and placed into the acid-paired context on day 4, females exhibited less hindpaw sensitivity than males (*p* = 0.011; **Fig. 2D**); male hindpaw withdrawal thresholds were lower in the acid-paired chamber on day 1 as compared to day 4 (*p* = 0.002), whereas female hindpaw withdrawal thresholds were higher in the acid-paired context on day 4 relative to day 1 (*p* = 0.006). This latter effect is especially interesting considering the dose of acetic acid administered in the acid-paired context on day 4 is 9-fold higher than the dose administered in this same context on day 1. Collectively, these data suggest that that prior experience, or pain memories associated with a specific environment, induce a conditioned compensatory response that decreases pain sensitivity – or in other words, increases pain tolerance – only in female mice.

### Female and male mice develop context-dependent pain tolerance after training with high doses of acetic acid

These initial experiments raised the possibility that males were not able to develop conditional pain tolerance. However, unlike females, male mice also failed to develop mechanical hypersensitivity (*i*.*e*., ‘anti-DNIC’) after injection of 0.1% acetic acid on day 1 of training **(Fig. 2B)**. Therefore, we repeated this paradigm but administered 0.9% acetic acid on all training and test days **(Fig. 3A)** to determine if the unconditional stimulus used in the previous experiments was simply not strong enough to support associative learning in males.

**Figure 3:**
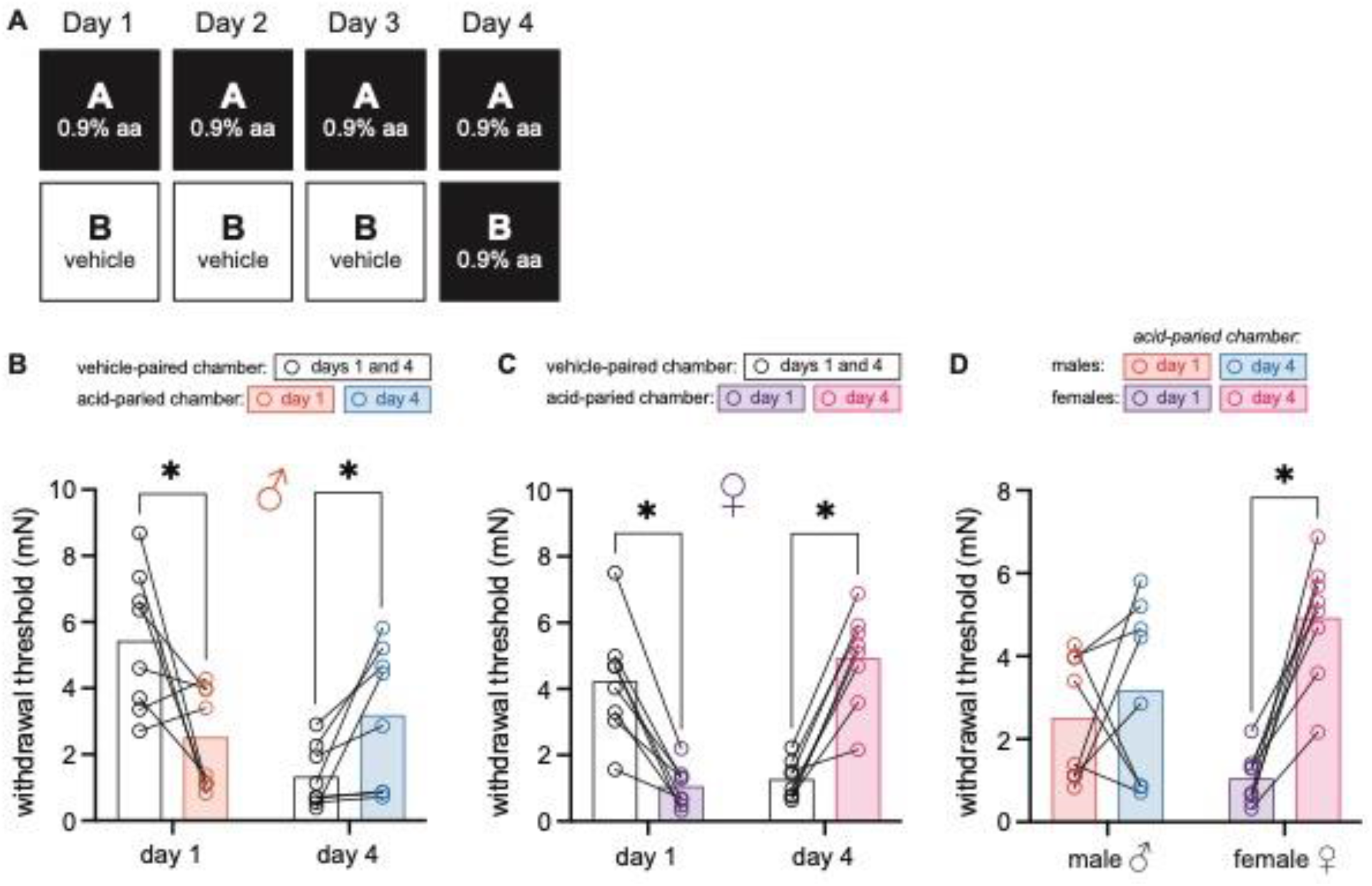
Female and male mice develop context-dependent pain tolerance after training with high doses of acetic acid. **A)** Experimental design depicting the within-subject procedure. In daily sessions, mice were given a 0.9% dose of acetic acid in one physical chamber, Context A, and vehicle injections in a separate physical chamber, Context B. On the final day, mice received an injection of 0.9% acetic acid solution in each context. Hindpaw sensitivity was measured 45 min following injection on day 1 and day 4. **B)** von Frey withdrawal thresholds of male mice (n=8) on day 1 and day 4 of the paradigm. Acetic acid injection on day 1 (0.9%) induced hindpaw mechanical hypersensitivity (‘anti-DNIC’). Hindpaw mechanical sensitivity differed between the contexts following 0.9% acetic acid injection on day 4, however; males exhibited contextually mediated pain tolerance (i.e., less pain sensitivity) in the acid-paired chamber relative to the vehicle-paired chamber on day 4. **C)** von Frey withdrawal thresholds of female mice (n=8) on day 1 and day 4 of the paradigm. Acetic acid injection on day 1 (0.1%) induced hindpaw mechanical hypersensitivity (‘anti-DNIC’). Hindpaw mechanical sensitivity differed between the contexts following 0.9% acetic acid injection on day 4, however; females exhibited contextually mediated pain tolerance (i.e., less pain sensitivity) in the acid-paired chamber relative to the vehicle-paired chamber on day 4. **D)** von Frey withdrawal thresholds of male and female mice replotted to highlight results in the acetic-acid paired chamber. Male mice exhibit similar levels of hindpaw mechanical sensitivity in the acid-paired chamber on days 1 and 4. Female mice exhibit less hindpaw sensitivity following 0.9% acetic acid injection on day 4 than they did following 0.9% acetic acid injection on day 1 suggesting the development of conditional pain tolerance.

The same 2 (Sex: Male, Female) x 2 (Context: Vehicle-Paired, Acetic Acid-Paired) x 2 (Day: 1, 4) ANOVA conducted to assess mechanical pain sensitivity found a marginal effect of day, *F*_(1, 14)_ = 4.12, MSE = 1.54, *p* = 0.06, ηp2 = 0.23, a day by sex interaction, *F*_(1, 14)_ = 12.26, MSE = 1.54, *p* = 0.004, ηp^2^ = 0.47, and a context by day interaction, *F*_(1, 14)_ = 40.63, MSE = 3.31, *p* < 0.001, ηp^2^ = 0.74. No other main effects or interactions were significant (largest *F* = 1.95, *p* = 0.12). As before, planned comparisons were conducted to assess differences both between- and within-groups.

In contrast to the results from the previous experiment, males exhibited hindpaw mechanical hypersensitivity in the acid-paired chamber on day 1 (*p* = 0.006; **Fig. 3B)** and conditional pain tolerance in this chamber on day 4 (*p* = 0.006) relative to the vehicle-paired chamber. A similar pattern was observed in females **(Fig. 3C)**; relative to the vehicle-paired chamber, female mice exhibited hindpaw mechanical hypersensitivity in the acid-paired chamber on day 1 (*p* = 0.003), and conditional pain tolerance in this chamber on day 4 (*p* < 0.001). Although the magnitude of ‘anti-DNIC’ exhibited following 0.9% acetic acid injection on day 1 was higher in females than males (*p* = 0.024; **Fig. 3D)**, the level of conditional pain tolerance exhibited in the acid-paired chamber on day 4 was not significantly different between the sexes (*p* = 0.078). In planned comparisons that assessed behavior across days, males exhibited ‘anti-DNIC’ in the vehicle-paired context on day 4 relative to day 1 (*p* < 0.001) but did not exhibit conditional pain tolerance in the acid-paired chamber on day 4 relative to day 1 (**Fig. 2D**; *p* = 0.42). This was not the case for females as they demonstrated both ‘anti-DNIC’ in the vehicle-paired context on day 4 relative to day 1 (*p* = 0.002) and conditional pain tolerance in the acid-paired context on day 4 relative to day 1 (*p* < 0.001). Considering these data, and the fact that male mice had higher withdrawal thresholds in the acid-paired context on day 4 as compared to the vehicle-paired context, we conclude that both sexes are able to develop context-dependent pain tolerance if trained with strong unconditional stimuli, but the magnitude of this phenomenon may differ between the sexes with females being especially sensitive to this type of learning.

### Conditional pain tolerance is not mediated by changes in circulating corticosterone, but rather increased endogenous opioid signaling

To begin probing the biological basis of this conditional pain tolerance, we next developed a between-subject 3-day conditioning paradigm in which animals were only trained in one environment so as to isolate physiological changes occurring in each context **(Fig. 4A)**. A 2 (Group: acid-trained, vehicle-trained) x 2 (Day: 1, 4) ANOVA found a day by group interaction, *F*_(1, 14)_ = 49.30, MSE = 2.01, *p* < 0.001, ηp^2^ = .78, but no main effect of group or day, largest *F* = 1.43, *p* = 0.25. Planned comparisons found that similar to observations in the acid-paired context in our within-subject experiment, acid-trained animals exhibited referred mechanical hypersensitivity relative to vehicle-trained animals on day 1 (*p* < 0.001) and conditional pain tolerance relative to vehicle-trained animals on day 4 (*p* = 0.008; **Fig. 4B)**. To determine if this context-dependent pain tolerance is a stress-mediated phenomenon, circulating corticosterone was measured in animals immediately upon removal from the testing environment. Corticosterone levels were similar between acid- and vehicle-trained mice **(Fig. 4C;** *t*_(14)_ = 0.56, *p* = 0.58) suggesting that the observed behaviors were not forms of stress-induced analgesia or hyperalgesia in the acid- and vehicle-trained mice respectively.

**Figure 4.**
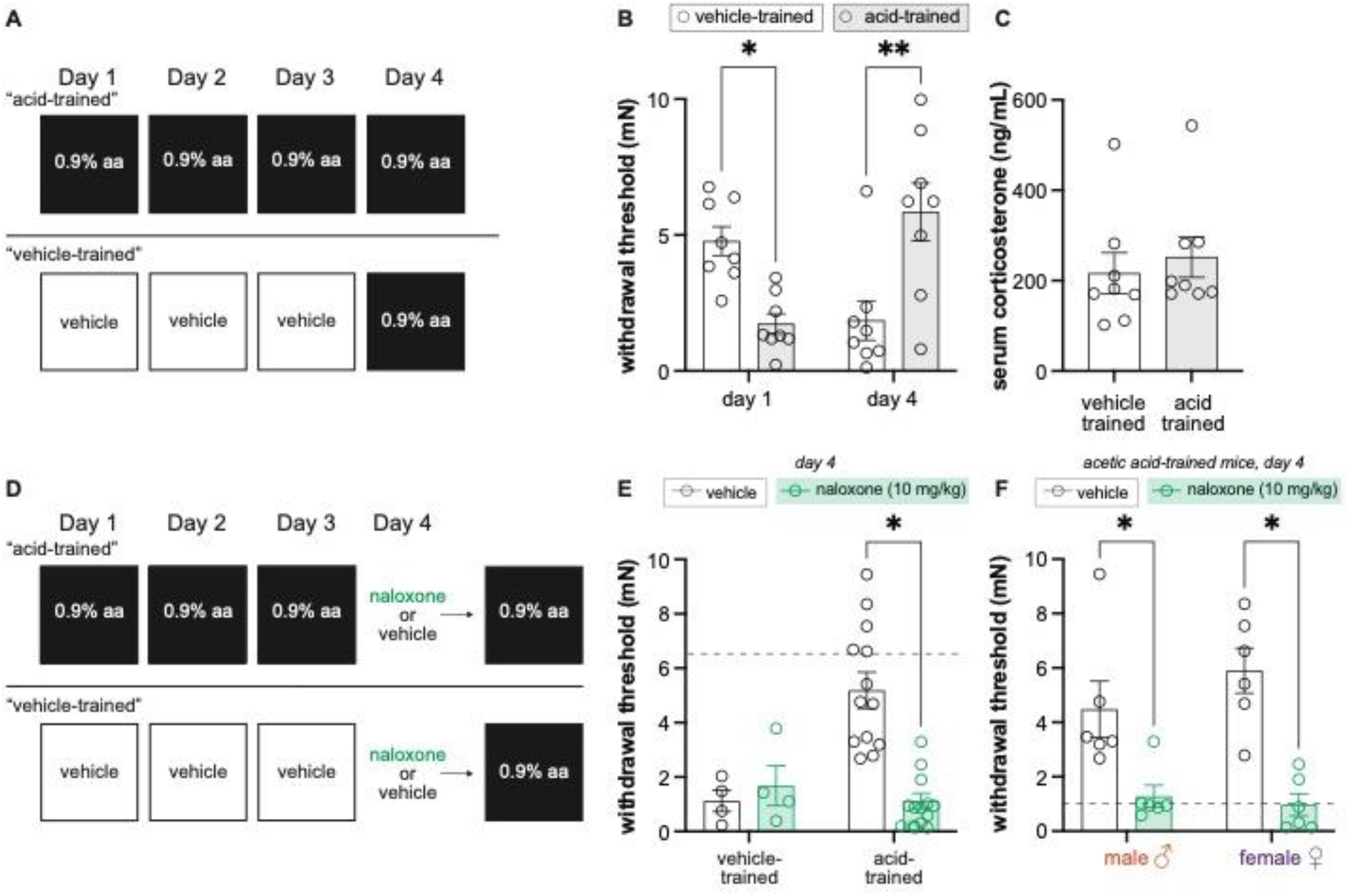
Endogenous opioid signaling, and not circulating corticosterone, is associated with contextually mediated conditional pain tolerance. **A)** Experimental design depicting the between-subjects design. Animals received an IP injection either acetic acid or vehicle for three days. On the fourth and final day, all animals were given an IP injection of acetic acid. **B)** von Frey withdrawal thresholds of male and female mice on day 1 and day 4 of the paradigm. Animals in the vehicle-trained group (n=8) have higher withdrawal thresholds than those injected with acid on day 1. On day 4, vehicle-trained mice (n=8) exhibit ‘anti-DNIC’ and have lower withdrawal thresholds than acid-trained mice. Acid-trained mice exhibit ‘anti-DNIC’ on day 1 and conditional pain tolerance on day 4. **C)** Results from ELISA corticosterone assay. Groups (n=8; same mice from panel B) did not differ in their circulating cortisol levels, suggesting that contextually mediated pain tolerance is not influenced by stress. **D)** Experimental design depicting the between-subjects design. Animals received an IP injection either acetic acid or vehicle for three days. On the fourth and final day, all animals were given an IP injection of acetic acid. However, half of the animals in each condition were pretreated with naloxone, an opioid antagonist, while the other half received vehicle. **E)** von Frey withdrawal thresholds of male and female mice on day 4. Animals in the vehicle-trained groups (n=4) showed mechanical hypersensitivity irrespective of naloxone pretreatment, and animals in the acid-trained group (n=12) exhibited decreased mechanical sensitivity that was blocked by naloxone. The dashed gray line represents the average withdrawal threshold of vehicle-trained mice on day 1. **F)** von Frey withdrawal thresholds of acid-trained male (n=6) and female (n=6) mice replotted to highlight sex differences on day 4. Both males and females showed increases in mechanical hypersensitivity when pre-treated with naloxone, suggesting that contextually mediated pain tolerance is mediated by the endogenous opioid system. The dashed gray line indicates the average withdrawal threshold of vehicle-trained mice on day 4.

Another possible biological explanation for conditional pain tolerance is that after repeated exposure to acetic acid, the context may promote the release of endogenous opioids as part of a compensatory response to the upcoming acetic acid treatment. To test this hypothesis, we repeated the between-subject 3-day conditioning paradigm, but injected animals with naloxone, a broad-spectrum opioid receptor antagonist, or vehicle before they were placed into the testing environment on day 4 **(Fig. 4D)**. A 2 (Group: vehicle-trained, acid-trained) x 2 (Drug: vehicle, naloxone) ANOVA found a main effect of group, *F*_(1, 28)_ = 6.68, MSE = 2.77, *p* = 0.015, ηp^2^ = 0.19, a main effect of drug, *F*_(1, 28)_ = 6.67, MSE = 2.77, *p* = 0.015, ηp^2^ = 0.19, and an interaction between the two, *F*_(1, 28)_ = 11.57, MSE = 2.77, *p* = 0.002, ηp^2^ = 0.29. Naloxone pre-treatment had no effect on the referred mechanical hypersensitivity exhibited by vehicle-trained animals on day 4 (*p* = 0.64), but it successfully blocked the conditional pain tolerance exhibited by acid-trained mice on day 4 **(Fig. 4E;** *p* < 0.001); the mechanical withdrawal thresholds of naloxone injected acid-trained mice were similar to those exhibited by vehicle-trained mice after their first exposure to acetic acid. To determine if naloxone treatment blocked conditional pain tolerance to a similar extent in acid-trained males and females, we performed a 2 (Sex: male, female) x 2 (Drug: vehicle, naloxone) ANOVA. This analysis revealed a main effect of drug, *F*_(1, 20)_ = 31.62, MSE = 3.14, *p* < 0.001, ηp^2^ = 0.61, but no main effect of sex nor an interaction between the two (*F*s < 1), suggesting that naloxone blocked conditional pain tolerance, in both male (vehicle vs. naloxone; *p* = 0.005) and female (vehicle vs. naloxone; *p <* 0.001) acid-trained mice **(Fig. 4F)**.

### Contextual cues previously associated with pain induce conditional hypersensitivity in male mice and conditional analgesia in female mice

Up to this point, all experimental paradigms included acid injection (UCS) on the final testing day. As a result, it was unclear if the compensatory analgesic response was truly context-dependent (*i*.*e*., a conditional response) or simply an autonomic response initiated by repeated acetic acid treatment. To answer this question, we developed a within-subject, 3-day training paradigm that concluded with vehicle administration before placement in either training context on day 4 **(Fig. 5A)**. von Frey withdrawal thresholds were analyzed using a 2 (Sex: Male, Female) x 2 (Context: Vehicle-Paired, Acetic Acid-Paired) x 2 (Day: 1, 4) ANOVA. This analysis found a main effect of context, *F*_(1, 14)_ = 46.45, MSE = 1.10, *p* < 0.001, ηp^2^ = 0.77, a day by sex interaction, *F*_(1, 14)_ = 4.79, MSE = 4.30, *p* = 0.046, ηp^2^ = 0.26, and a context by day interaction, *F*_(1, 14)_ = 5.65, MSE = 4.78, *p* = 0.032, ηp^2^ = 0.29, but no other main effects or interactions (largest *F* = 2.89, *p* = 0.11).

**Figure 5.**
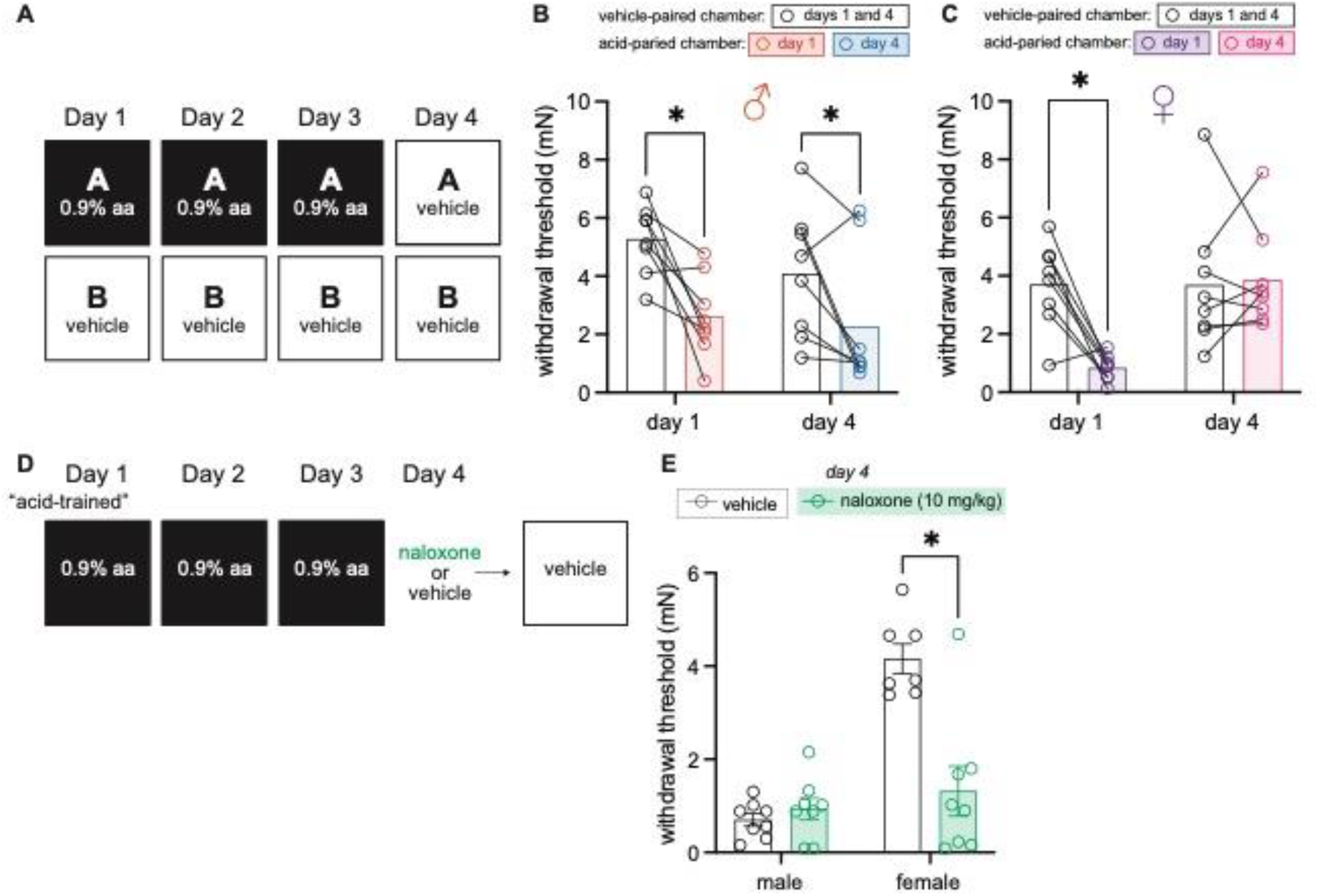
Males exhibit conditional hypersensitivity and females exhibit conditional analgesia in the absence of UCS administration. **A)** Experimental design depicting the within-subject procedure. In daily sessions, mice were given a 0.9% dose of acetic acid in one physical chamber, Context A, and vehicle injections in a separate physical chamber, Context B. On the final day, mice received an injection of vehicle in each context. Hindpaw sensitivity was measured 45 min following injection on day 1 and day 4. **B)** von Frey withdrawal thresholds of male mice (n=8) on day 1 and day 4 of the paradigm. Acetic acid injection on day 1 (0.9%) induced hindpaw mechanical hypersensitivity (‘anti-DNIC’). Similar hindpaw mechanical hypersensitivity was observed in the acid-paired chamber on day 4, despite the fact that animals received a vehicle injection prior to being placed in this chamber. **C)** von Frey withdrawal thresholds of female mice (n=8) on day 1 and day 4 of the paradigm. Acetic acid injection on day 1 (0.9%) induced hindpaw mechanical hypersensitivity (‘anti-DNIC’). Hindpaw mechanical sensitivity did not differ between the contexts following vehicle injection on day 4. **D)** Experimental design. In daily sessions, mice were given a 0.9% dose of acetic acid in one physical chamber. On the final day, mice received an injection of naloxone or vehicle prior to being placed into the same training context; no acid injection was performed on day 4. E) von Frey withdrawal thresholds of male (n=8) and female (n=7-8) mice on day 4 of the paradigm. Naloxone pre-treatment did not affect the hindpaw withdrawal thresholds of acid-trained male mice. Conversely, naloxone treatment decreased the hindpaw withdrawal thresholds of acid-trained female mice, suggesting conditional recruitment of endogenous opioid systems in the training context.

Male mice exhibited ‘anti-DNIC’ (*i*.*e*., hindpaw mechanical hypersensitivity in the acid-paired context relative to the vehicle-paired context; *p*<0.001; **Fig. 5B)** on day 1, and, similar to the conditional hypersensitivity previously reported after single exposure to a noxious UCS (Martin et al., 2019), context-dependent hypersensitivity in the absence of the UCS on day 4 (*p* = 0.027). Male hindpaw withdrawal thresholds in the acid-paired chamber were lower than thresholds recorded in the vehicle-paired chamber on day 4, despite receiving vehicle injections before being placed into either chamber. This conditional hypersensitivity was not observed in females. Despite exhibiting the same ‘anti-DNIC’ effect as males on day 1 (*p* < 0.001; **Fig. 5C)**, female mice exhibited the same level of mechanical sensitivity in the vehicle- and acid-paired contexts on day 4 **(Fig. 5C**, *p* = 0.81). Further, in contrast to male mice that exhibited similar withdrawal thresholds in the acid-paired chamber on day 1 and day 4, female hindpaw withdrawal thresholds in the acid-paired chamber were significantly higher on day 4 than on day 1 (*p* = 0.003), suggesting either a lack of any conditional response, or the development of conditional analgesia.

To determine if female behavior on day 4 was reflective of context-dependent endogenous opioid release, and thus a form of conditional analgesia, we performed a between-subject, 3-day acetic acid training paradigm and treated animals with naloxone prior to being placed in the testing context on day 4 **(Fig. 5D)**. Mechanical sensitivity was assessed on day 4 **(Fig. 5E)** using a 2 (Sex: male, female) x 2 (Drug: vehicle, naloxone) ANOVA. This found main effects of both sex, F_(1,27)_ = 31.42, MSE = .91, *p* < 0.001, ηp^2^ = 0.54, drug, F_(1,27)_ = 14.44, MSE = .91, *p* < 0.001, ηp^2^ = 0.35, and an interaction between the two, F_(1,27)_ = 20.07, MSE = .91, *p* < 0.001, ηp^2^ = 0.43. Naloxone treatment had no effect on the conditional hypersensitivity exhibited by male mice on day 4 (*p* = 0.63), further supporting the notion that this behavior is an example of stress-induced hyperalgesia. When administered in females however, naloxone unmasked an endogenous opioid-mediated conditional compensatory response (*p* < 0.001). In the absence of naloxone treatment, female behaviors appear to be unaffected by 3 days of acid-training (*i*.*e*., females had high withdrawal thresholds in the acid-trained context on day 4 relative to day 1 as they are not receiving an acetic acid injection prior to entering the context; **Fig. 5C)**. However, in the absence of acetic acid injection on day 4, naloxone treatment decreased withdrawal thresholds suggesting that females exhibit context-dependent conditional hypersensitivity that they readily counteract with activation of their endogenous opioid system.

## Discussion

In the present experiments we asked whether environment can exert control over pain sensitivity and pain expectation through associative learning. We first demonstrated, in a within-subject manner, that a context paired with a painful experience (acetic acid injection; UCS) can control conditional pain tolerance. Although female mice demonstrate this conditional pain tolerance after pairing a relatively weak UCS with the CS, male mice only develop conditional pain tolerance if a stronger UCS is used throughout conditioning trials. We then determined that this effect is not mediated by differences in circulating corticosterone, but rather by activation of the endogenous opioid system since administration of the opioid antagonist naloxone blocked conditional pain tolerance in both sexes. In a second set of experiments, we demonstrated that males exhibit conditional hypersensitivity when placed into a context previously associated with pain. Alternatively, females exhibit conditional analgesia that is dependent on endogenous opioid signaling when placed into a similar environment. Together, these results demonstrate that context can come to exert control over the endogenous opioid system such that it is recruited as a conditional compensatory response to ongoing painful stimuli in both sexes, or in anticipation of a painful stimulus in females. Further, these results are the first (to our knowledge) to demonstrate that acute visceral pain can be used as an unconditional stimulus to elicit conditional analgesia; previous explorations in conditional analgesia used footshock, a noxious somatic stimuli that also induces freezing behaviors.

In the ongoing search for better/novel analgesics, many preclinical biologists, including some of the present authors, have taken a reductionist approach using highly standardized *in vitro* or *in vivo* assays to determine how a gene or protein contributes to the process of nociception. In so doing, significantly less has been learned about the influence external factors have on pain perception and the development of chronic pain. In their ‘Imprecision Hypothesis’ (2015), Moseley and Vlaeyen posit that chronic pain may develop, in part, as a result of associative learning; repeated coupling of a UCS– in this case a noxious input – with inert cues like a specific movement of the body (*e*.*g*., bending over) or context, may lead to the perception of pain (CR) upon subsequent presentation of these newly conditioned stimuli. Data presented in this manuscript are, to our knowledge, the first preclinical test of this hypothesis; we repeatedly paired noxious visceral stimulation (*i*.*e*., intraperitoneal acetic acid injection; UCS) with a unique context (*i*.*e*., a behavior chamber; CS). Similar to the results obtained following a single pairing of this same UCS and CS (Martin et al., 2019), male mice developed conditional hypersensitivity upon CS presentation alone **(Fig. 5B)**. In contrast, when females are exposed to the CS, they engage their endogenous opioid system, such that behavioral responses appear similar to those exhibited prior to injury or conditioning **(Fig. 5C)**. These divergent results suggest there are qualitative sex differences in the biological processes that mediate pain expectation.

The biological basis and evolutionary explanation for these qualitative sex differences is currently unknown. Dimorphisms in endogenous opioid circuit anatomy and function are well documented (Chakrabarti et al., 2010; Liu et al., 2007; Loyd et al., 2006; 2008; Tershner et al., 2000). Perhaps pressures associated with the reproduction timeframe and energy requirement led to differential abilities to recruit endogenous opioid signaling such that female organisms are better able to activate this system as a means of dampening ongoing pain. Another intriguing hypothesis is that brain regions required for contextual conditioning – like the dorsal hippocampus (Maren & Holt, 2003; Matus-Amat et al., 2004) and retrosplenial cortex (Keene & Bucci, 2008; Kwapis et al., 2015; Miller et al., 2021; Trask et al., 2021)—and those involved in descending pain inhibition are differentially connected in each sex. For example, electrical stimulation of the retrosplenial cortex was shown to decrease formalin-associated spontaneous pain (Reis et al., 2010) and the mechanical hypersensitivity that develops following a surgical incision in male rats (Rossaneis et al., 2011). Notably however, no female rodents were included in these studies. Regardless, this electrically evoked analgesia required opioid signaling in the anterior pretectal nucleus (Rossaneis et al., 2011), a brain region that receives direct input from the retrosplenial cortex. Future studies should examine (1) if this circuit functions similarly in female rodents, (2) if this circuit is engaged during contextual conditioning, and (3) if the retrosplenial cortex, or other structures important for contextual learning, differentially projects to other brain regions associated with endogenous analgesia modulation in the two sexes.

In addition to assessing pain expectation, we also examined if associative learning impacts pain sensitivity, or the magnitude of the UCR elicited by a noxious stimulus. The UCS used in these experiments was intraperitoneal injection of acetic acid, a procedure that induces reflexive writhing behaviors (Koster et al., 1959). The UCR used in these experiments was hindpaw mechanical sensitivity testing. Hindpaw hypersensitivity can develop in visceral pain models through a phenomenon known as referred pain (Bielefeldt et al., 2006; Lamb et al., 2006), the biological basis of which is traditionally ascribed to C fiber axonal bifurcation (*i*.*e*., one axonal branch detects and relays somatic information while another detects and relays visceral information) or convergence of viscero-somatic information at second order spinal cord neurons. More recently however, Tansley et al. (2019) characterized a phenomenon in which pain in one part of the body causes hyperalgesia in an anatomically separate body region if evoked with a relatively innocuous stimulus. In contrast to the ‘pain-inhibits-pain’ idea associated with diffuse noxious inhibitory control (DNIC), this observation, termed ‘facilitatory conditioned pain modulation (CPM)’ or ‘anti-DNIC’, is observed across multiple rodent species and mouse strains and does not depend on the anatomical proximity of the injured and test sites. Despite initially demonstrating referred pain/’anti-DNIC’, both male and female mice in the present experiments developed conditional pain tolerance after repeated exposure to the noxious stimuli; both sexes exhibited less pain when presented with a noxious UCS in an environment (CS) associated with an acute pain memory. Notably, however, there was a quantitative sex difference in the strength of the UCS required to develop this conditional pain tolerance; male mice only developed conditional pain tolerance if trained with high doses of acetic acid. This is perhaps unsurprising given the extensive literature describing sex differences in pain perception.

The idea that males and females perceive pain differently is a long-standing topic in both academic and popular science discussions. Upon review of this topic, it is clear that women are more sensitive to acute pain stimuli than men, and that chronic pain is more frequently reported in women than it is in men (Mogil, 2012; Mogil, 2020). Rodent data presented in this paper are in agreement with this conclusion; female mice exhibited more pain than male mice following low (0.1%; **Fig. 2D)** and high (0.9%; **Fig. 3D)** acetic acid administration. Similar quantitative sex differences have been previously reported in other visceral pain models. For example, male and ovariectomized female rodents are less sensitive to chemical or mechanical injury of the colon and urinary bladder (reviewed by Traub and Ji 2014). Combined with the fact that stronger UCS will support more conditioning to stimuli presented with those UCS (Holland, 1979; Morris & Bouton, 2006; see Rescorla & Wagner, 1972), the present results suggest that females more readily associate contextual cues with noxious visceral stimuli because they perceive visceral pain more intensely than males. Under this logic, females should develop conditional pain tolerance more quickly than males. In support of this hypothesis, 6 of 8 female mice included in **Figure 3** exhibited conditional pain tolerance on day 2 of the paradigm (*i*.*e*., higher paw withdrawal thresholds in the acid-paired chamber on day 2 vs. day 1) whereas 0 of the 8 male mice in this same study exhibited conditional pain tolerance on day 2. Further, recall that females developed conditional pain tolerance when lower acetic acid concentrations were used as the UCS while males did not **(Fig. 2)**. When applied to a diverse patient population, this hypothesis suggests that females, and particularly those with heightened pain sensitivities, may be more susceptible to partake in associative learning with painful unconditional stimuli. Although opportunity for maladaptive learning exists, the inverse of this may also be true; females may be better prepared, biologically speaking, to associate relief of ongoing pain with conditional stimuli than males. Confirmation of this hypothesis may lead to the development of novel cognitive therapeutic strategies for pain that rely on the principles of Pavlovian conditioning, or those related to other memory-based manipulations like reframing (Pavlova et al., in press).

In conclusion, the data presented here demonstrate that context can control pain perception while an animal is actively experiencing pain or when an animal is anticipating a painful experience. These results are evidence that pain perception and engagement of endogenous opioid systems can become dependent on the environment for expression and can be modified through their psychological association with environmental cues. Future work will examine the neural pathways associated with the contextual control of pain processing including those that mediate opioid release in response to pain-associated environments. Further, additional studies will need to examine if continuous chronic pain is susceptible to the effects of conditioning observed here using procedures that result in acute, transient pain.

## References

Bielefeldt, K., Lamb, K., & Gebhart, G. F. (2006). Convergence of sensory pathways in the development of somatic and visceral hypersensitivity. American Journal of Physiology-Gastrointestinal and Liver Physiology, 291(4), G658–G665.

Bouton, M. E., & Bolles, R. C. (1979). Contextual control of the extinction of conditioned fear. Learning and Motivation, 10(4), 445–466.

Chakrabarti, S., Liu, N. J., & Gintzler, A. R. (2010). Formation of μ-/κ-opioid receptor heterodimer is sex-dependent and mediates female-specific opioid analgesia. Proceedings of the National Academy of Sciences, 107(46), 20115–20119.

Chaplan, S. R., Bach, F. W., Pogrel, J. W., Chung, J. M., & Yaksh, T. L. (1994). Quantitative assessment of tactile allodynia in the rat paw. Journal of Neuroscience Methods, 53(1), 55–63.

Chesler, E. J., Ritchie, J., Kokayeff, A., Lariviere, W. R., Wilson, S. G., & Mogil, J. S. (2003). Genotype-dependence of gabapentin and pregabalin sensitivity: the pharmacogenetic mediation of analgesia is specific to the type of pain being inhibited. Pain, 106(3), 325–335.

Dixon, W. J. (1980). Efficient analysis of experimental observations. Annual Review of Pharmacology and Toxicology, 20(1), 441–462.

Dixon, W. J. (1965). The up-and-down method for small samples. Journal of the American Statistical Association, 60(312), 967–978.

Koster R, Anderson M, De Beer E. 1959. Acetic acid for analgesic screening. Fed Proc, 18, 412–430.

Fanselow, M. S., & Helmstetter, F. J. (1988). Conditional analgesia, defensive freezing, and benzodiazepines. Behavioral Neuroscience, 102(2), 233.

Helmstetter, F. J., & Fanselow, M. S. (1987). Effects of naltrexone on learning and performance of conditional fear-induced freezing and opioid analgesia. Physiology & Behavior, 39(4), 501–505.

Holland, P. C. (1979). The effects of qualitative and quantitative variation in the US on individual components of Pavlovian appetitive conditioned behavior in rats. Animal Learning & Behavior, 7, 424–432.

Keene, C. S., & Bucci, D. J. (2008). Neurotoxic lesions of retrosplenial cortex disrupt signaled and unsignaled contextual fear conditioning. Behavioral Neuroscience, 122(5), 1070.

Koster, R., Anerson, M, & De Beer, E. J. (1959). Acetic acid for analgesic screening. Federation Proceedigs (Vol. 18, p. 412).

Kwapis, J. L., Jarome, T. J., Lee, J. L., & Helmstetter, F. J. (2015). The retrosplenial cortex is involved in the formation of memory for context and trace fear conditioning. Neurobiology of Learning and Memory, 123, 110–116.

Lamb, K., Zhong, F., Gebhart, G. F., & Bielefeldt, K. (2006). Experimental colitis in mice and sensitization of converging visceral and somatic afferent pathways. American Journal of Physiology-Gastrointestinal and Liver Physiology, 290(3), G451–G457.

Lee, G. J., Kim, S. A., Kim, Y. J., & Oh, S. B. (2021). Naloxone-induced analgesia mediated by central kappa opioid system in chronic inflammatory pain. Brain Research, 1762, 147445.

Liu, N. J., von Gizycki, H., & Gintzler, A. R. (2007). Sexually dimorphic recruitment of spinal opioid analgesic pathways by the spinal application of morphine. Journal of Pharmacology and Experimental Therapeutics, 322(2), 654–660.

Long, C. C., Sadler, K. E., & Kolber, B. J. (2016). Hormonal and molecular effects of restraint stress on formalin-induced pain-like behavior in male and female mice. Physiology & Behavior, 165, 278–285.

Loyd, D. R., & Murphy, A. Z. (2006). Sex differences in the anatomical and functional organization of the periaqueductal gray-rostral ventromedial medullary pathway in the rat: A potential circuit mediating the sexually dimorphic actions of morphine. Journal of Comparative Neurology, 496(5), 723–738.

Loyd, D. R., Wang, X., & Murphy, A. Z. (2008). Sex differences in μ-opioid receptor expression in the rat midbrain periaqueductal gray are essential for eliciting sex differences in morphine analgesia. Journal of Neuroscience, 28(52), 14007–14017.

Maren, S., & Holt, W. G. (2004). Hippocampus and Pavlovian fear conditioning in rats: muscimol infusions into the ventral, but not dorsal, hippocampus impair the acquisition of conditional freezing to an auditory conditional stimulus. Behavioral neuroscience, 118(1), 97.

Martin, L. J., Acland, E. L., Cho, C., Gandhi, W., Chen, D., Corley, E., … & Mogil, J. S. (2019). Male-specific conditioned pain hypersensitivity in mice and humans. Current Biology, 29(2), 192–201.

Matus-Amat, P., Higgins, E. A., Barrientos, R. M., & Rudy, J. W. (2004). The role of the dorsal hippocampus in the acquisition and retrieval of context memory representations. Journal of Neuroscience, 24(10), 2431–2439.

Miller, A. M., Serrichio, A. C., & Smith, D. M. (2021). Dual-Factor Representation of the Environmental Context in the Retrosplenial Cortex. Cerebral Cortex, 31(5), 2720–2728.

Mogil, J. S. (2020). Qualitative sex differences in pain processing: emerging evidence of a biased literature. Nature Reviews Neuroscience, 21(7), 353–365.

Mogil, J. S. (2012). Sex differences in pain and pain inhibition: multiple explanations of a controversial phenomenon. Nature Reviews Neuroscience, 13(12), 859–866.

Morris, R. W., & Bouton, M. E. (2006). Effect of unconditioned stimulus magnitude on the emergence of conditioned responding. Journal of Experimental Psychology: Animal Behavior Processes, 32(4), 371.

Moseley, G. L., & Vlaeyen, J. W. (2015). Beyond nociception: the imprecision hypothesis of chronic pain. Pain, 156(1), 35–38.

Pavlov, I. P. (1927). Conditioned reflexes: an investigation of the physiological activity of the cerebral cortex.

Pavlova, M., Lund, T., Nania, C., Kennedy, M., Graham, S., & Noel, M. (in press). Reframe the pain: A randomized controlled trial of a parent-led memory-reframing intervention. The Journal of Pain.

Reis, G. M., Dias, Q. M., Silveira, J. W. S., Del Vecchio, F., Garcia-Cairasco, N., & Prado, W. A. (2010). Antinociceptive effect of stimulating the occipital or retrosplenial cortex in rats. The Journal of Pain, 11(10), 1015–1026.

Rescorla, R. A., & Wagner, A. R. (1972). A theory of Pavlovian conditioning: Variations in the effectiveness of reinforcement and nonreinforcement. In A.H. Black & W. F. Prokasy (Eds.), Classical Conditioning II: Current Research and Theory (pp. 64–99). New York, NY: Appleton-Century-Crofts.

Rosen, S. F., Ham, B., Haichin, M., Walters, I. C., Tohyama, S., Sotocinal, S. G., & Mogil, J. S. (2019). Increased pain sensitivity and decreased opioid analgesia in T-cell-deficient mice and implications for sex differences. Pain, 160(2), 358–366.

Rossaneis, A. C., Reis, G. M., & Prado, W. A. (2011). Stimulation of the occipital or retrosplenial cortex reduces incision pain in rats. Pharmacology Biochemistry and Behavior, 100(2), 220–227.

Siegel, S. (1977). Morphine tolerance acquisition as an associative process. Journal of Experimental Psychology: Animal Behavior Processes, 3(1), 1.

Siegel, S., Hinson, R. E., Krank, M. D., & McCully, J. (1982). Heroin” overdose” death: contribution of drug-associated environmental cues. Science, 216(4544), 436–437.

Stevenson, G. W., Bilsky, E. J., & Negus, S. S. (2006). Targeting pain-suppressed behaviors in preclinical assays of pain and analgesia: effects of morphine on acetic acid-suppressed feeding in C57BL/6J mice. The Journal of Pain, 7(6), 408–416.

Tansley, S. N., Macintyre, L. C., Diamond, L., Sotocinal, S. G., George, N., Meluban, L., … & Mogil, J. S. (2019). Conditioned pain modulation in rodents can feature hyperalgesia or hypoalgesia depending on test stimulus intensity. Pain, 160(4), 784–792.

Tershner, S. A., Mitchell, J. M., & Fields, H. L. (2000). Brainstem pain modulating circuitry is sexually dimorphic with respect to mu and kappa opioid receptor function. Pain, 85(1-2), 153–159.

Trask, S., & Bouton, M. E. (2014). Contextual control of operant behavior: evidence for hierarchical associations in instrumental learning. Learning & Behavior, 42(3), 281–288.

Trask, S., Pullins, S. E., Ferrara, N. C., & Helmstetter, F. J. (2021). The anterior retrosplenial cortex encodes event-related information and the posterior retrosplenial cortex encodes context-related information during memory formation. Neuropsychopharmacology, 46(7), 1386–1392.

Trask, S., Reis, D. S., Ferrara, N. C., & Helmstetter, F. J. (2020). Decreased cued fear discrimination learning in female rats as a function of estrous phase. Learning & Memory, 27(6), 254–257.

Traub, R. and Ji, Y. (2014). Sex differences and hormonal modulation of deep tissue pain. Front Neuroendocrinol, 34(4).

Watkins, L. R., Wiertelak, E. P., & Maier, S. F. (1993). The amygdala is necessary for the expression of conditioned but not unconditioned analgesia. Behavioral Neuroscience, 107(2), 402.

